# Elucidating the Differential Impacts of Equivalent Gating-Charge Mutations in Voltage-Gated Sodium Channels

**DOI:** 10.1101/2024.09.09.612021

**Authors:** Eslam Elhanafy, Amin Akbari Ahangar, Rebecca Roth, Tamer M. Gamal El-Din, John R Bankston, Jing Li

## Abstract

Voltage-gated sodium (Na_v_) channels are pivotal for cellular signaling and mutations in Na_v_ channels can lead to excitability disorders in cardiac, muscular, and neural tissues. A major cluster of pathological mutations localizes in the voltage-sensing domains (VSDs), resulting in either gain-of-function (GoF), loss-of-function (LoF) effects, or both. However, the mechanism behind this functional divergence of mutations at equivalent positions remains elusive. Through hotspot analysis, we identified three gating charges (R1, R2, and R3) as major mutational hotspots in VSDs. The same amino-acid substitutions at equivalent gating-charge positions in VSD_I_ and VSD_II_ of the cardiac sodium channel Na_v_1.5 show differential gating-property impacts in electrophysiology measurements. We conducted 120 µs molecular dynamics (MD) simulations on wild-type and six mutants to elucidate the structural basis of their differential impacts. Our μs-scale MD simulations with applied external electric fields captured VSD state transitions and revealed the differential structural dynamics between equivalent R-to-Q mutants. Notably, we observed transient leaky conformations in some mutants during structural transitions, offering a detailed structural explanation for gating-pore currents. Our salt-bridge network analysis uncovered VSD-specific and state-dependent interactions among gating charges, countercharges, and lipids. This detailed analysis elucidated how mutations disrupt critical electrostatic interactions, thereby altering VSD permeability and modulating gating properties. By demonstrating the crucial importance of considering the specific structural context of each mutation, our study represents a significant leap forward in understanding structure-function relationships in Na_v_ channels. Our work establishes a robust framework for future investigations into the molecular basis of ion channel-related disorders.

## Introduction

As one of the most widely distributed types of ion channels, voltage-gated sodium (Na_v_) channels initiate action potentials and serve a central role in electrical excitability by selectively allowing sodium (Na^+^) ions to flow through the cell membrane (Catterall, 2010; Hille, 1987). Heartbeats, muscle twitches, and lightning-fast thoughts are all manifestations of the bioelectrical signals that rely on the activity of Na_v_ channels. More than 2000 mutations in human Na_v_ channels have been associated with various heart, muscle, and brain excitability disorders (George, 2005; Ghovanloo et al., 2016; Huang et al., 2017; Pan et al., 2018). For instance, disruption of the activation and inactivation of the cardiac Na_v_1.5 channel has been identified as a major cause of long QT syndrome type 3 (LQT3), Brugada syndrome type 1 (BRGDA1) (Kapplinger et al., 2010; Millat et al., 2006), and other arrhythmias (Li et al., 2018; Remme et al., 2008; Remme and Bezzina, 2010; Ruan et al., 2009). However, this genetic discovery raises important mechanistic questions about how similar mutations can cause opposing functional changes, such as gain-of-function (GoF) and loss-of-function (LoF). Addressing these questions is crucial for developing selective therapeutics for precision medicine.

The Na_v_ channel’s α subunit exhibits a heterotetrametric structure, with each of the four repeats encompassing six transmembrane (TM) helices (S1-S6). The pore domain (PD) is composed of S5, S6, and the pore loop (P-loop) from all repeats, while the voltage-sensing domain (VSD) is formed by the first four TMs (S1 to S4) (Catterall, 2000). The structures of the Na_v_ channel consistently show that the structural transition of VSD is mediated by the perpendicular sliding of the S4 helix through an hourglass-shaped structure formed by the S1, S2, and S3 segments of the VSD (Catterall et al., 2020; Clairfeuille et al., 2017).

The VSDs, are essential for sensing membrane potential and initiating sodium channel activation/recovery and inactivation (Catterall, 2010), contain the S4 helix with four to six gating charges (arginine or lysine) arrayed across the membrane as the voltage sensor (Fig. 1) (Yarov-Yarovoy et al., 2012). These gating charged residues (R1 to R6) are counterbalanced by negatively charged residues in the S1–S3 helices, which are also named countercharges (Fig. 1). Both gating charge and countercharge residues are highly conserved across different isoforms (Groome and Bayless-Edwards, 2020). A conserved aromatic residue in S2, known as the hydrophobic constriction site (HCS), acts as a steric barrier to S4 translocation (Pless et al., 2014; Schwaiger et al., 2013). The countercharges above the HCS from each helix are situated in an extracellular negatively charged cluster (ENC) and are referred to as S1E, S2E, and S3E. Similarly, those closer to the cytoplasmic region occupy the intracellular negatively charged cluster (INC) and are referred to as S1I, S2I, and S3I (Fig. 1). Interactions of countercharges with gating charges have been investigated in a set of functional experiments that support roles for countercharges in channel activation and S4 translocation (Andrew M. Glazer et al., 2024; Groome and Bayless-Edwards, 2020; Moreau et al., 2014a; Pless et al., 2014; Shen et al., 2024).

**Figure 1.**
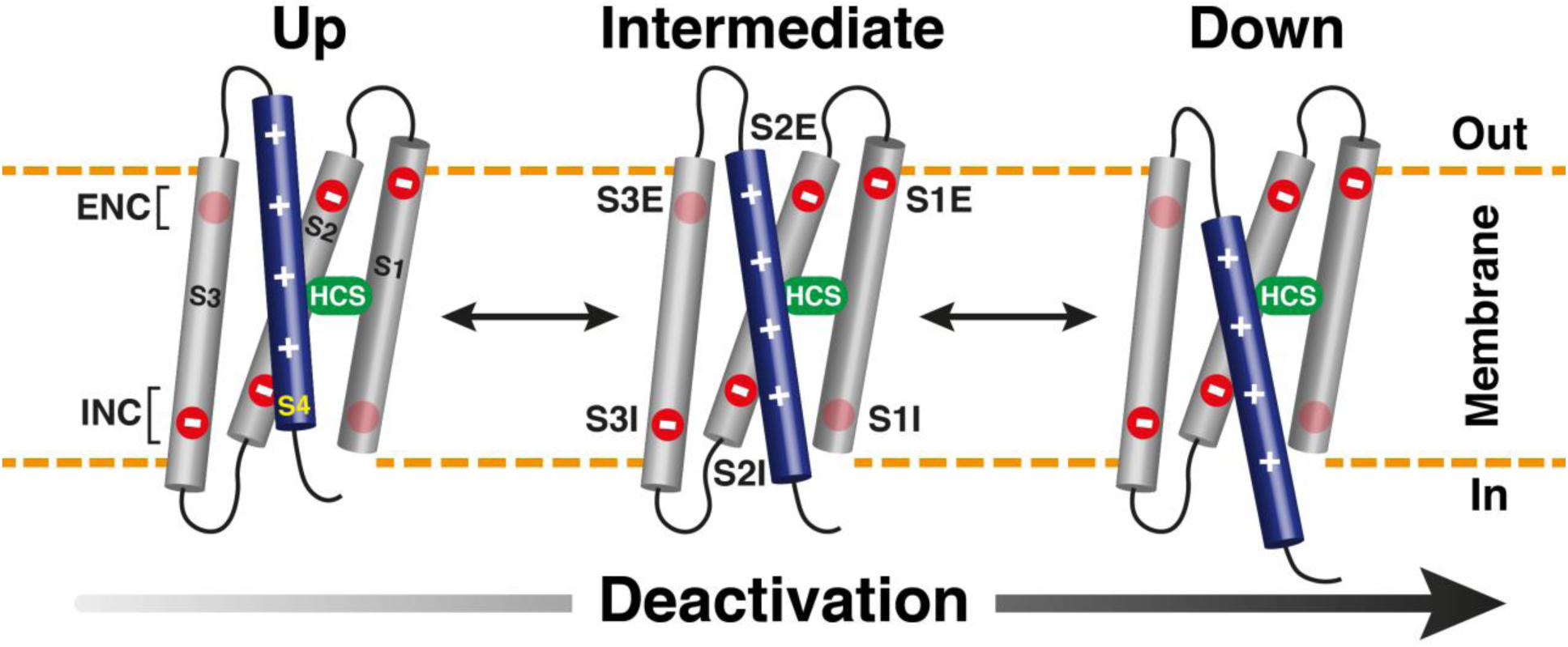
Stepwise deactivation of VSDs of Na_v_ channels. Cartoon depiction of the sequential deactivation of the Na_v_1.5 VSD in response to membrane potential changes. The S4 helix (blue) moves relative to static S1-S3 (gray), with negatively (-) charged countercharged residues (red) and the hydrophobic constriction site (HCS) residue (green). The S4 positively (+) charged gating charge residues adopt an up conformation under depolarized potential and transition to a down conformation as the potential recovers, illustrating the VSD’s dynamic nature.

Current models suggest that the gating charges of S4 traverse an aqueous gating pore within the VSD (Schwaiger et al., 2013). Under resting membrane potential, the S4 gating charges adopt a down conformation (Fig. 1). The depolarization of membrane potential drives the S4 to an up conformation, activating the channel. Mutations of the S4 gating charges disrupt interactions with countercharges (Moreau et al., 2015b, 2014a, 2014b), creating a new permeation pathway known as the gating pore (Gosselin-Badaroudine et al., 2012a; Sokolov et al., 2007, 2005; Struyk and Cannon, 2007; Tombola et al., 2005). Gating pores represent an alternative permeation pathway that emerges within the typically non-conductive VSDs of voltage-gated ion channels (Gosselin-Badaroudine et al., 2012a; Jiang et al., 2018; Moreau et al., 2015b, 2015a; Sokolov et al., 2007; Struyk et al., 2008). Gating-pore currents, also known as omega currents (I_ω_), have been suggested as a common pathological mechanism linking various mutations occurring in the VSDs of Na_v_ channels (Eltokhi et al., 2024; Jiang et al., 2018; Moreau et al., 2015a, 2015b; Struyk et al., 2008).

VSD mutations showed diverse impacts across various gating properties, such as maximum current amplitude (I_Max_), I_ω_, time constant of recovery from inactivation (τ_rec_), and voltage at half-maximal activation (V_1/2act_) and inactivation (V_1/2inact_) (Ahangar et al., 2024). These changes result in overall GoF, LoF, or mixed effects, contributing to the complexity of Na_v_ channelopathies. However, the mechanisms underlying why and how equivalent gating-charge mutations produce different impacts on gating properties and diverse functional phenotypes remain unclear. To address this critical question, it is necessary to perform a comprehensive structural analysis of each mutation, especially considering the intricate interactions between the mutation site and its surrounding structural environment. More importantly, ion channel function is not only influenced by static structures but also deeply rooted in their transitions among multiple functional states. Therefore, the effects of mutations are determined by the intricate interplay between structural elements and their coordinated movements during functional transitions. Mutations can potentially alter these structural dynamics, ultimately affecting gating properties and channel function. The complexity of ion channel structures and the dynamic nature of their functional transitions necessitate a biophysical approach with atomic resolution and dynamic description to understand the molecular mechanisms underlying various mutational impacts. Molecular dynamics (MD) is such an ideal approach for this study and the availability of cryo-EM structures (Huang et al., 2022a; Jiang et al., 2021, 2020; Li et al., 2022, 2021; Pan et al., 2021, 2019, 2018; Shen et al., 2019) also provides a unique opportunity to address this fundamental question using MD simulations.

To systematically compare the differential impacts of gating charge mutations, we focused on R-to-Q mutations at the R1 to R3 positions in VSD_I_ and VSD_II_ of the cardiac sodium channel Na_v_1.5. A total of 120 µs of MD simulations were conducted, including three independent runs for the WT and six R-to-Q mutants in VSD_I_ and VSD_II_. The MD simulations were coupled with appropriate external electric fields to accelerate VSD structural transitions at µs timescale. Based on MD trajectories of WT and six mutants, a detailed analysis, particularly a state-dependent salt-bridge network analysis, was performed to reveal how each mutation distinctly affects the structural transitions of the VSDs.

## Materials and methods

### Model and simulation systems building

The Cryo-EM structure of human Na_v_1.5 (PDB: 7DTC) (Li et al., 2021) is used for constructing the VSD_I_ and VSD_II_ systems. Each system encompassed two segments: the voltage-sensing domain (VSD) and the pore domain (PD). For constructing the system of VSD_I_, not only the VSD of D_I_ (residues 119 to 250) but also the PD of D_II_ (residues 842 to 944) were included to maintain a native VSD-PD interface. PD of D_II_ is restrained through simulations. Considering only the PD of D_II_ is included and thus the polar selectivity filter residues in PD_II_ face lipids. To prevent unfavorable contacts between hydrophilic selectivity filter residues and hydrophobic lipid tails, targeted mutations to Alanine were introduced in three residues within the pore loop of D_II_. Mirroring the approach used for VSD_I_, the VSD_II_ system comprised two segments: the VSD of D_II_ (residues 699 to 838) and the PD of D_III_ (residues 1329 to 1480). Similar to VSD_I_, mutations to Alanine were implemented in three residues of D_III_. Additionally, disulfide bonds were assigned according to the information provided in the PDB structure (Li et al., 2021). To ascertain the correct protonation states of ionizable residues, pKa calculations were conducted employing PROPKA3 (Olsson et al., 2011; Søndergaard et al., 2011). This resulted in a model where all residues maintained their default protonation states. Subsequently, the protein’s first principal axis was aligned with the z-axis using the OPM (Orientations of Proteins in Membranes) database (Lomize et al., 2012). The lipid composition of the heart exhibited a high abundance of POPC (1-palmitoyl-2-oleoyl-sn-glycero-3-phosphocholine) and POPI (1-palmitoyl-2-oleoyl-sn-glycero-3-phosphoinositol) compared to other tissues (Pradas et al., 2018; Tomczyk and Dolinsky, 2020). To observe native lipid-protein interactions, the finalized systems were then embedded into a bilayer composed of POPC and POPI in a 3:1 ratio. A system area of (80 x 80 Å^2^), was constructed with the membrane’s normal aligned along the z-axis using the CHARMM-GUI Membrane builder (Wu et al., 2014). Then, the systems were hydrated by introducing a 10 Å layer of water to each side of the membrane. Finally, the system total charge was neutralized with a 150 mM NaCl solution. The ultimate dimensions of the system before equilibration were 80 × 80 × 85 Å^3^, comprising approximately 51,000 atoms.

### Molecular dynamics simulations

Simulations described in this work were performed using NAMD software (Phillips et al., 2005) (version 2.14 or 3.0) or Desmond (Bowers et al., 2006) on the specialized computational platform Anton2 (Shaw et al., 2014). We used CHARMM36 (Huang and Mackerell, 2013) parameters for the protein (Best et al., 2012; Huang et al., 2016; Huang and Mackerell, 2013) and lipids (Klauda et al., 2010), respectively, along with the TIP3P model for explicit water molecules (Jorgensen et al., 1983) and the associated ionic parameters with NBFIX corrections (Luo and Roux, 2010; Noskov and Roux, 2008; Venable et al., 2013). All simulations were performed under tetragonal periodic boundary conditions (PBCs) to the simulation box to overcome finite-size effects and mimic bulk-like properties. The simulations were performed with a time step of 2 fs. Throughout the simulations, all covalent bonds involving hydrogen atoms were constrained using the SHAKE (Ryckaert et al., 1977) algorithm. Electrostatic and van der Waals interactions were computed at each simulation step for maximum accuracy.

Following 5,000 steps of energy minimization, all systems were simulated using the following protocol: (i) 1ns constant pressure and constant temperature (NPT) simulation with all heavy atoms constrained, (ii) 1ns NPT simulation with all carbon α atoms constrained, and (iii) Equilibration in an NPT ensemble with PD (residues 842 to 944 in D_II_ and residues 1329 to 1480 in D_III_) constrained to reach 30ns allowing proper hydration of solvent-exposed regions of the Na_v_ pore cavity using NAMD. For MD simulations using NAMD, the system was simulated in the NPT ensemble using the Nosé−Hoover Langevin piston method to maintain the pressure at 1 atm and a Langevin thermostat to maintain the temperature at 310 K (Feller et al., 1995; Martyna et al., 1994). The oscillation period of the piston was set at 100 fs and the damping time scale at 50 fs. Long-range electrostatic interactions were calculated using the particle mesh Ewald (PME) algorithm (Darden et al., 1993). Short-range non-bonded interactions were calculated with a cutoff of 12 Å and the application of a smoothing decay started to take effect at 10 Å.

After the initial equilibration, the systems were subjected to production simulations in the NPT ensemble using Desmond on Anton2 (Supplementary Table 1). For MD simulations conducted using Desmond on Anton2, a Berendsen coupling scheme was implemented to sustain a consistent pressure of 1.0 atm. The calculation of long-range electrostatic interactions was facilitated by the k-space Gaussian split Ewald method (Shan et al., 2005). All MD trajectories were visualized and analyzed using VMD,(Humphrey et al., 1996) in-house Tcl, and Python scripts.

### Analysis

<u>The z-position distance analysis</u> was used to track the movement of gating charged residues, namely R219, R222, R225, and R228 in VSD_I_, and R808, R811, R814, R817, and K820 in VSD_II_. This analysis focuses on gating charge movement relative to HCS residues Tyr168 in VSD_I_ and Phe760 in VSD_II_, delineating the boundary between the extracellular and intracellular hydrated regions of the VSDs. This analysis measures the distance along the z-direction between the center of mass of the side chain of each gating-charge residue and the center of mass of HCS. The z-position distance analysis was conducted using a combination of in-house Tcl script and Python script for data visualization. The Tcl script was used for calculations in VMD to analyze the MD trajectories, while the Python script was programmed for data visualization. This integrated approach allows for continuous monitoring of each gating charge’s position throughout the simulation period, precisely measuring the transition rate for each gating charge.

<u>HOLE Analysis</u> (Smart et al., 1996) with an updated interface from MDAnalysis (Gowers et al., 2016; Michaud-Agrawal et al., 2011) was employed to analyze and visualize the gating-pore radius within VSDs throughout the simulation trajectories. By focusing on the conformational difference caused by the movement of gating-charge residues, HOLE quantified the size of the aqueous pathway that can traverse the transmembrane pore of the VSD. Based on Monte Carlo simulated annealing, the algorithm identifies optimal routes for a sphere with a variable radius to pass through the channel. In the plot, only the minimum radius along the pore is selected for each frame and plotted as a function of time. Specifically, S1-S4 protein segments from VSD_I_ (residues 131 to 230) and VSD_II_ (residues 717 to 819) were selected for HOLE analysis to avoid non-native hole detections. To validate the pore surface, we cross-checked it using VMD. Our software pipeline involves HOLE calculations and Python scripts for data visualization.

<u>Salt Bridge Network Analysis</u> investigates salt bridges formed within a protein throughout a simulation trajectory. Specifically, we focused on gating charges (R219, R222, R225, and R228 in VSD_I_, and R808, R811, R814, R817, and K820 in VSD_II_) and countercharge (D152, E161, E171, D197 in VSD_I_, and D720, E737, E763, E785, and E795 in VSD_II_) residues. Our analysis included the following steps: We first identified gating charges, countercharge residues, and phosphate head groups from lipid molecules. We then measured the distances between the terminal carbons of the side chains of countercharge (acidic) residues and gating charge (basic) residues. The distances between phosphate head groups of lipids and the terminal carbon of gating-charged residues were measured similarly. Subsequently, the output data was analyzed using a Python script to calculate the occupancy percentage of the salt-bridge interaction at each state, with a cutoff distance of 5 Å to define the formation of a salt bridge. Additionally, transitions were determined based on z-position distance analysis to separate distinct phases for each state (up, intermediate, down) for each trajectory. Occupancy was calculated by dividing the number of frames with a distance ≤ 5 Å over the total number of frames in each state following the equation.

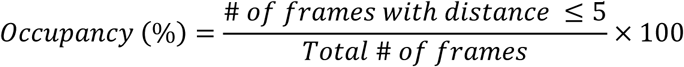

<u>Ion Permeation Analysis</u> tracks Na^+^ and Cl^-^ ions permeability through VSD during simulation trajectories. This method calculates the distance between each ion in a 4 Å radius of HCS residue in each VSD (Y168 in VSD_I_ and F760 in VSD_II_). It then plots the relative positions of all ions along the z coordinate, connecting consecutive points less than 20 Å apart. This approach allowed for a detailed examination of ion permeations through VSDs.

### Supplementary Materials

It provides detailed experimental electrophysiology data, including methods for electrophysiological recordings and data analysis of both WT and mutant systems. This section is supported by 2 figures and 2 tables. Supplementary Figure 1 presents functional measurements for R2 and R3 to glutamine mutations in VDS_I_ and VSD_II_. Supplementary Figure 2 illustrates the time series of VSD structural transitions, gating pore openings, and ion permeation events. Supplementary Table 1 summarizes the MD system composition, simulation durations, and the number of replica simulations for WT and mutant systems. Supplementary Table 2 provides a summary of the gating properties of the mutations. This comprehensive data set enhances understanding of the studied mutations’ electrophysiological characteristics and structural dynamics.

## Results

### Gating charges are major mutational hotspots with diverse functional impacts

In our recent study (Ahangar et al., 2024), over 2,400 disease-associated missense variants annotated in the UniProt database (Famiglietti et al., 2019; McGarvey et al., 2019) were mapped across nine human Na_v_ channels to search the most representative pathological variants. The VSD is identified as a major cluster of mutation hotspots. Mutations in the VSD exhibit diverse impacts on gating properties, including I_max_, τ_rec_, V_1/2 Act_, and V_1/2 Inact_ with no clear preference between GoF and LoF effects (Ahangar et al., 2024). The disease-associated missense variants annotated from VSDs are mapped onto the multiple sequence alignment (MSA) shown in (Fig. 2). Notably, a high prevalence of mutation hotspots is observed within the S4 helix, specifically located at gating charges (Fig. 3). For instance, there are 26, 17, and 26 mutations located on R1, R2, and R3 of all VSDs (Fig. 3A), respectively, which is significantly higher than the number of mutations found in other residues within the Na_v_ channel. In Na_v_1.5 VSD_I_, R219 is associated with 5 phenotypes, R222 with 12 phenotypes, and R225 with 9 phenotypes (Fig. 3B). Similarly, in VSD_II_, R808 is linked to 8 phenotypes, R811 to 5 phenotypes, and R814 to 8 phenotypes (Fig. 3C). Another mutation hotspot is observed in the S3 segment of VSDs. This site is identified as having a highly conserved countercharge residue (S3I), which plays a crucial role in interacting with gating charges to influence structural transitions. The specific residues at S3I are D197 in VSD_I_ and D785 in VSD_II_. These findings highlight the importance of gating-charge and countercharge residues in the function of Na_v_ channels.

**Figure 2.**
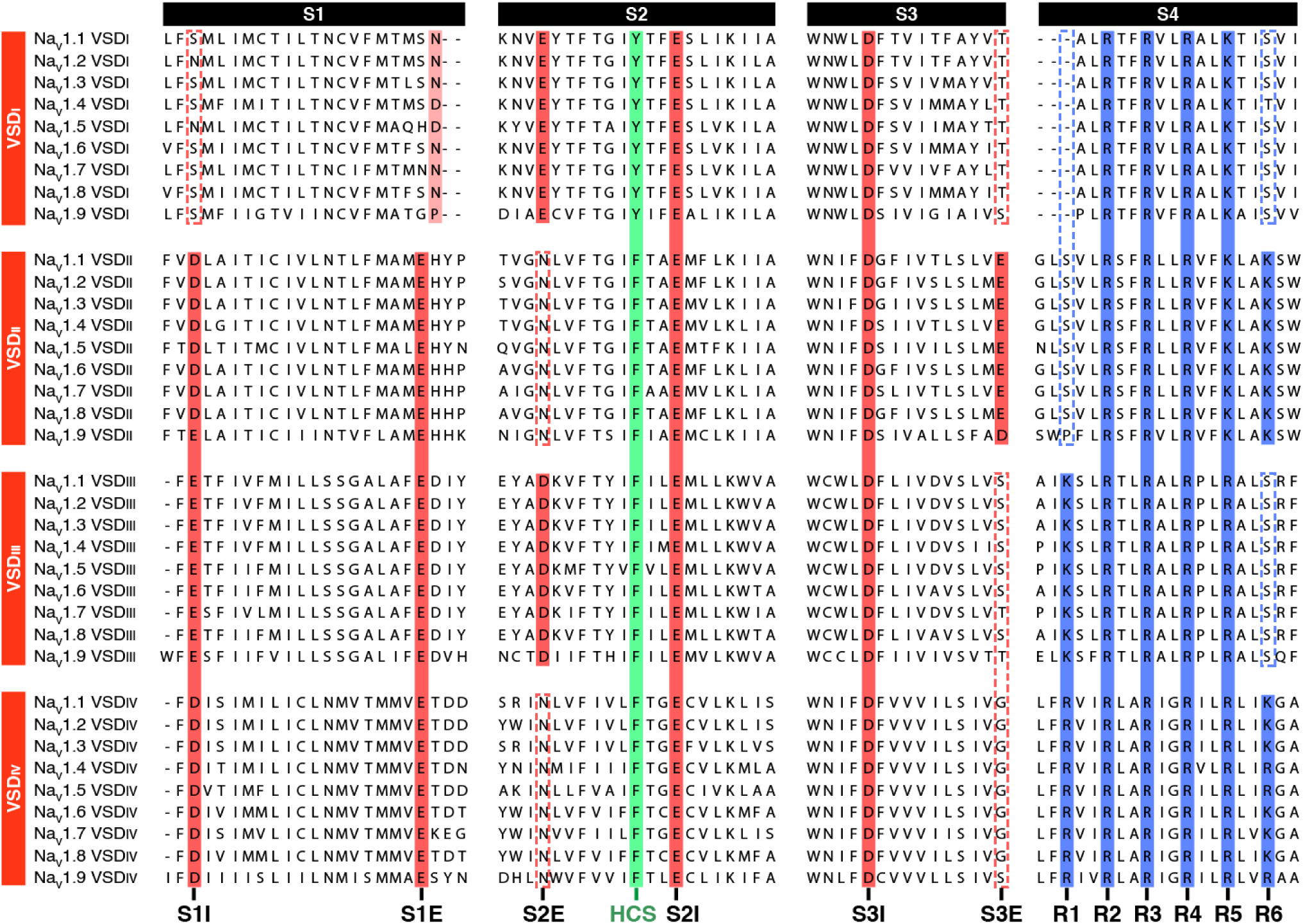
**Multiple sequence alignment of VSDs from four repeats of nine isoforms in the human Na_v_ channel family**, highlighting countercharge residues (red), HCS (green), and gating charge residues (blue).

**Figure 3.**
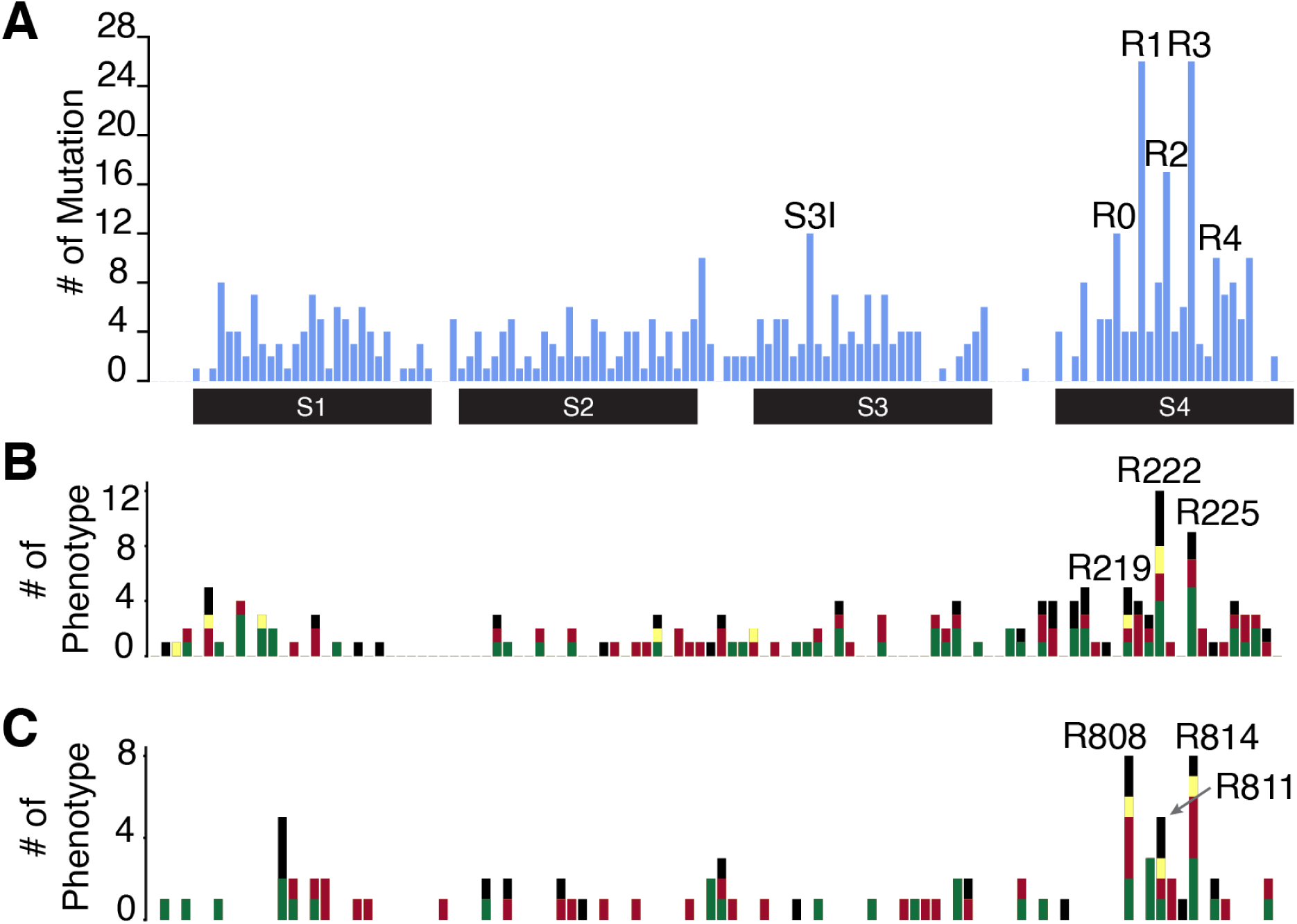
Mapping the mutation hotspots in VSDs of Na_v_ channels. The annotated disease- associated mutations from UniProt are mapped along the multiple sequence alignment (MSA) of VSDs in 9 human Na_v_ channels. The number of mutations at the equivalent position in 4 VSDs (A), the number of phenotypes in VSD_I_ (B), and that in VSD_II_ (C) are used to demonstrate the mutation hotspots. (B, C) pathogenic missense mutations are colored according to GoF (green bars), LoF (red), mixed (yellow), and uncertain (black) effects. Hotspots are labeled in this figure with either the order of gating charges (A) or residue IDs from Na_v_1.5 (B, C).

Disease-associated mutations at the gating-charge residues disrupt the activity of VSDs and lead to diverse functional impacts. Based on previous literature, comparing disease associations reveals that equivalent gating-charge mutations in VSD_I_ and VSD_II_ of Na_v_1.5 result in different phenotypes (Table 1). Some of the observed differences can be attributed to specific amino acid substitutions. For example, R to Q and R to W replacements in the same position can lead to distinct outcomes. This is exemplified by the R814Q mutation resulting LQT3, whereas R814W causes BRGDA1 (Glazer et al., 2020; Moreau et al., 2015a; Nguyen et al., 2008).

**Table 1.** The documented disease associations and gating-property impacts of R1, R2, and R3 gating-charge mutations of VSD_I_ and VSD_II_.

| Mutation Residue ID in Na<sub>v</sub>1.5 |  | Disease-associated Phenotypes from Na<sub>v</sub>1.5 mutations † |  | Gating Properties for Na<sub>v</sub>1.5 | References |
| --- | --- | --- | --- | --- | --- |
| VSD<sub>I</sub> |  |  |  |  |  |
| R1 | R219 | Q | No identified disease association to date | No significant change | (Chen et al., 1996; Gosselin-Badaroudine et al., 2012b; Moreau et al., 2018) |
|  |  | H/P | PFHB1A, LQT3, CMD1E, BRGDA1 | I<sub>ω</sub>: ↑<br>V<sub>1/2act</sub>: →<br>V<sub>1/2inact</sub>: ← |  |
| R2 | R222 | Q | CMD1E, RWS, LQT3, BRGDA1 | I<sub>Max</sub>: ↑<br>I<sub>ω</sub>: ↑<br>V<sub>1/2act</sub>: ←<br>V<sub>1/2inact</sub>: ← | (Daniel et al., 2019; Laurent et al., 2012; Mann et al., 2012; Moreau et al., 2015b; Nair et al., 2012) |
| R3 | R225 | Q | LQT3 | V<sub>1/2act</sub>: →<br>V<sub>1/2inact</sub>: → | (Beckermann et al., 2014; Chen et al., 1996; Moreau et al., 2015a, 2015b; Strege et al., 2018) |
|  |  | W/P | PFHB1A, BRGDA1, CIAT | I<sub>Max</sub>: ↑ / ↓*<br>I<sub>p</sub>: ↑<br>I<sub>ω</sub>: ↑<br>V<sub>1/2act</sub>: ← / →*<br>τ<sub>rec</sub>: ↑ / ↓* |  |
| VSD<sub>II</sub> |  |  |  |  |  |
| R1 | R808 | Q | No identified disease association to date | V<sub>1/2act</sub>: →<br>V<sub>1/2inact</sub>: → | (Chen et al., 1996; Glazer et al., 2020) |
|  |  | H/C | BRGDA1, RWS | I<sub>Max</sub>: ↓<br>V<sub>1/2inact</sub>: ←<br>τ<sub>rec</sub>: ↓ |  |
| R2 | R811 | H | BRGDA1 | I<sub>Max</sub>: ↓<br>V<sub>1/2inact</sub>: ←<br>τ<sub>rec</sub>: ↓ | (Calloe et al., 2013) |
| R3 | R814 | Q | BRGDA1 | I<sub>p</sub>: ↑<br>V<sub>1/2inact</sub>: ← | (Chen et al., 1996; Glazer et al., 2020; Kapplinger et al., 2010; Moreau et al., 2015a; Nguyen et al., 2008) |
|  |  | W | CMD1E | I<sub>ω</sub>: ↑<br>V<sub>1/2act</sub>: ← / →*<br>τ<sub>rec</sub>: ↓ |  |
| →: right-shifted, ←: left-shifted, ↑: increase, ↓: decrease |  |  |  |  |  |

\* Indicates a discrepancy between multiple mutations or references for the same gating property. † Disease names are reported as OMIM IDs. The associated phenotype and gating-property impacts for R-to-Q mutations are highlighted in red. **PFHB1A**: Progressive Familial Heart Block, Type 1A, **LQT3**: Long QT syndrome 3, **CMD1E**: Dilated Cardiomyopathy 1E, **BRGDA1**: Brugada syndrome, **RWS**: Romano-Ward Syndrome.

To focus on the same amino-acid substitution at the equivalent positions, we investigated the effects of identical R-to-Q mutations in the R2 position of VSD_I_ and VSD_II_ in Na_v_1.5 channels under consistent conditions, electrophysiological measurements were conducted on R2 equivalent mutations (R222Q and R811Q). The results demonstrated that the same mutation at an equivalent position exhibits diverse effects on channel gating properties, including alterations in voltage-dependence of activation, inactivation, and recovery from inactivation (Supp. Fig. 1). Previous experimental measurements and molecular modeling studies also reveal distinctive impacts on gating properties caused by R3 equivalent mutations in Na_v_1.5 (Table 1). R225Q (R3 in VSD_I_) mutation is associated with GoF disease (LQT3) (Chen et al., 1996), while R814Q (R3 in VSD_II_) mutation is associated with LoF diseases (BRGDA1) (Glazer et al., 2020). A key functional difference between these mutants is the shift in V_1/2inact_, which is left-shifted in R814 mutants but right-shifted in R225 mutants (Chen et al., 1996; Glazer et al., 2020). Additionally, while neither R219Q nor R808Q were reported to be linked to specific channelopathies (Table 1), R808Q exhibited a rightward shift in both V_1/2act_ and V_1/2inact_, unlike R219Q, which showed no significant difference compared to WT (Chen et al., 1996). These differences underscore the complexity and VSD-specific effects of these mutations and suggest that the gating properties and disease associations of a specific mutation cannot be presumed to be similar to those of its equivalent mutation.

In this study, six mutants in R1 to R3 positions of Na_v_1.5, including R219 (R1 in VSD_I_), R808 (R1 in VSD_II_), R222 (R2 in VSD_I_), R811 (R2 in VSD_II_), R225 (R3 in VSD_I_), and R814 (R3 in VSD_II_), were selected for a systematic simulation investigation. The selection of these mutants provides a focused approach to understanding the differential impacts of equivalent gating-charge mutations.

### An external electric field triggers VSD state transition in simulation on the µs scale

To investigate the structural role of a mutation in functional transitions, we needed to develop a computational protocol that enabled us to characterize structural transitions on an accessible microseconds (μs) timescale in MD simulations. An intrinsic property of voltage-gated ion channels is that their gating behaviors are driven by the change in the membrane potential. Thus, MD simulation was adapted by coupling different electric fields to mimic depolarization and repolarization to study the structural transition of VSDs. After exploring different electric fields, ±500 mV has been determined as the appropriate voltage to trigger the structural transition within a few µs without evident protein unfolding.

Starting from the Cryo-EM structure of Na_v_1.5 in a fast-inactivated state where the VSD is in the up position (PDB ID: 7DTC)(Li et al., 2021), μs-scale MD simulations were conducted to investigate the state transition on Na_v_1.5 VSDs. A 13 μs simulation of VSD_II_ characterized the structural transitions of the VSD responding to different external electric fields. In the initial 3 μs under an external electric field of -500 mV, the S4 helix, which bears five gating-charge residues (R1, R2, R3, K4, and K5), exhibited ∼10 Å sliding along the z-direction toward the cytoplasm (Fig. 4), which is consistent with the proposed model based on known structures (Catterall et al., 2020; Clairfeuille et al., 2017; Huang et al., 2022b). During the first 3 μs, the initiation of the gating-charge movement is attributed to the R3 residue crossing the HCS at 1.3 μs (Fig. 4), which represents an intermediate between down and up states. Following this, the R2 residue crossed the HCS in turn at 2 μs, and inward S4 motion typically stopped at this down state when R1 is above the HCS and directly contacts F760. During this conformational change, the VSD transitioned from an up state with three gating charges above HCS to a down state with one gating charge above the HCS (Fig. 4). Subsequently, when the direction of the external electric field was reversed to +500 mV over the next 4 μs (3 – 7 μs), the gating charges reverted to the up state with three gating charges above the HCS. A subsequent reversal of the electric field to -500 mV led to the VSD reaching the down-minus state at 10 μs, with four gating charges below the HCS. Finally, when the external electric field was switched back to +500 mV after 10 μs, the VSD returned to the up state at 13 μs (Fig. 4).

**Figure 4.**
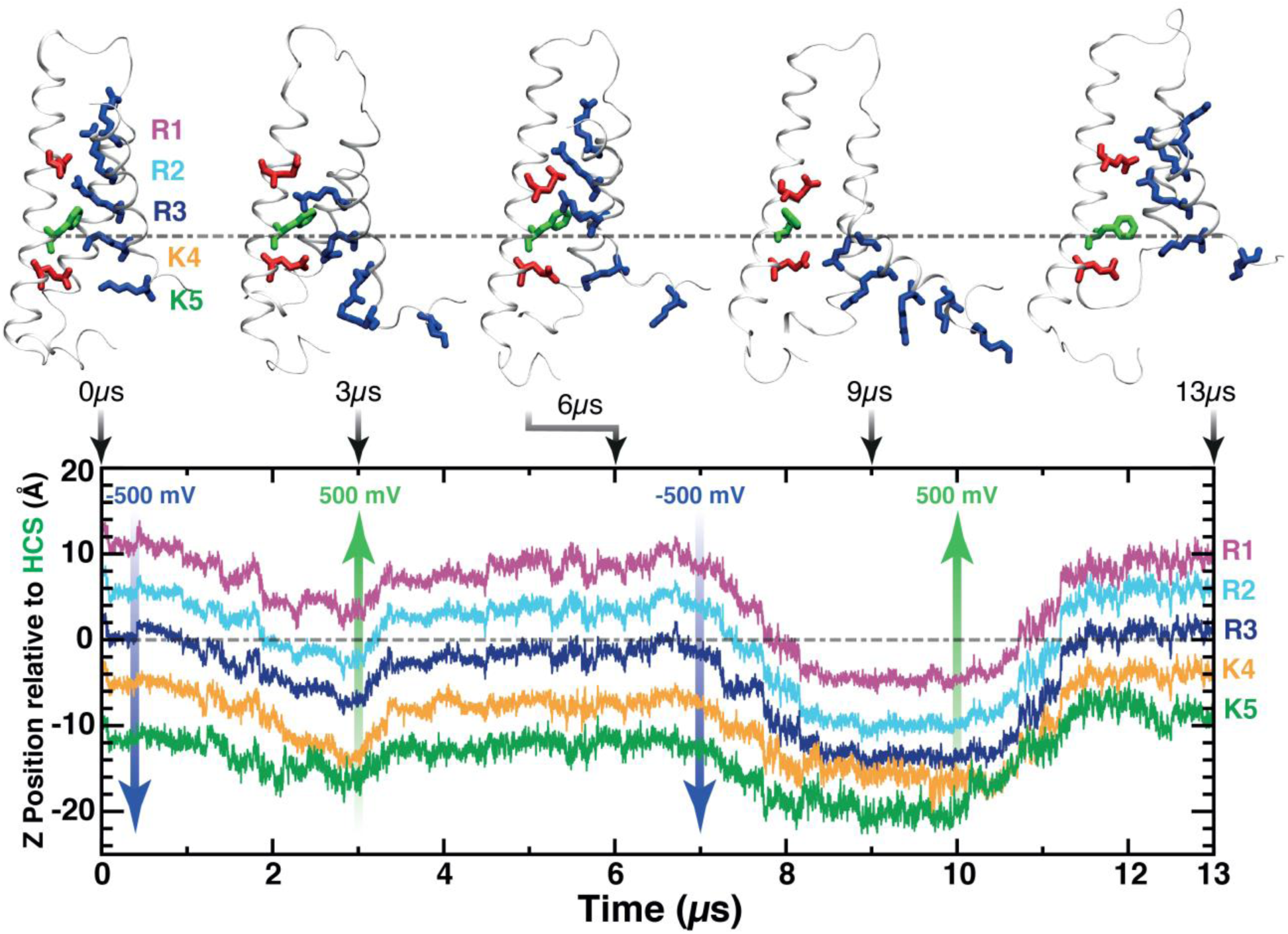
State transitions of VSD in a µs-scale MD simulation. The dynamic behavior of VSD_II_ under an external electric field of 500mV in opposite directions. Traces of z positions for Cα atoms of the gating charge residues (R1-K5) in different colors in VSD_II_ relative to the HCS are shown to track the structural changes. Molecular images of representative VSD_II_ snapshots (0, 3, 6, 9, and 13 µs) are presented in a white ribbon representation, with licorice representations of gating charges (blue), countercharges (red), and the HCS (green). The direction and magnitude of the external electric field applied are depicted by arrows (blue and green).

During the simulation trajectory, the gating charges shifted to contact new countercharges, forming electrostatic interactions. This adjustment was crucial for accommodating the gating charges to stabilize each structural state. Notably, no remarkable misfolding was observed in VSD_II_ during the simulation. This observation not only confirms the structural stability of VSDs under these high electric fields but also validates the feasibility of using this external electric field (±500 mV) to characterize the state transition of the VSDs within a few μs. This approach enables the simulation of conformational responses of the VSD (both WT and mutants) to depolarization and repolarization of membrane potential within the currently achievable simulation timescale. Accordingly, MD simulations were performed under the identical electric fields for both the WT and mutants within VSD_I_ (R219Q, R222Q, and R225Q) as well as VSD_II_ (R808Q, R811Q, and R814Q).

### Equivalent R-to-Q gating-charge mutations demonstrate differential structural dynamics

Under identical conditions and the same external electric field, simulations revealed that the equivalent mutations from R-to-Q in the gating charge of VSD_I_ and VSD_II_ exhibited differential structural dynamics toward the down state. All simulations were initiated from an up state, characterized by having three gating charges that are above HCS, under an applied external electric field of -500 mV. The z-position of gating charges was monitored to determine the structural states of mutants in both VSD_I_ and VSD_II_. As shown in (Fig. 5), distinct dynamic behaviors between equivalent R-to-Q mutants were observed.

**Figure 5.**
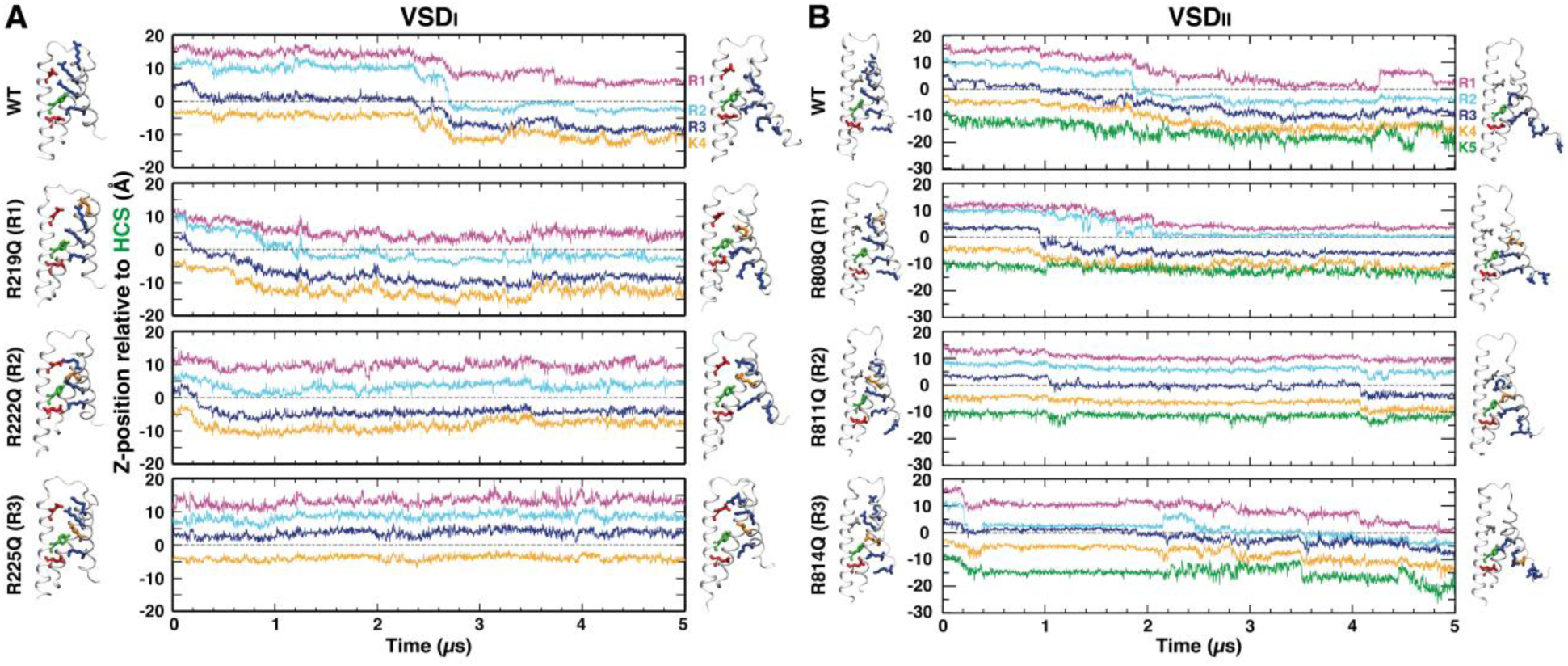
Differential impacts of R-to-Q mutations in VSD_I_ and VSD_II_ during up-to-down transitions. The differential dynamic behaviors of R-to-Q mutations (R1, R2, and R3) in VSD_I_ (A) and VSD_II_ (B) in comparison to the wild type (WT) under an external electric field of -500mV. Traces of z positions for Cα atoms of the gating-charge residues in S4 relative to that of the HCS are shown in each panel to track the structural changes. Molecular images of VSDs are presented in a white cartoon representation, illustrating the initial and final frames with licorice representations of gating charges (blue), countercharges (red), and the HCS (green). Only one representative trajectory among three independent runs for each mutant is illustrated here.

Upon comparing equivalent mutants from VSD_I_ and VSD_II_, distinct dynamic behaviors were observed. A typical example is the R3 mutants (R225Q in VSD_I_ and R814Q in VSD_II_) that exhibited highly differential structural dynamics under the same condition. In VSD_I_, the R3 mutant (R225Q) showed resistance to transition. Unlike the WT, it remained in the up state throughout the entire duration of the simulation time (Fig. 5A), this persistence was consistently observed in three independent runs for R225Q (Fig. 6A). However, in VSD_II_, the R3 mutant (R814Q) showed WT-like behavior as it exhibited a complete transition from the up state to the down state (Fig. 5B). This complete transition was also observed in another independent run of R814Q (Fig. 6B). Conversely, the R2 mutants (R222Q in VSD_I_ and R811Q in VSD_II_) exhibited distinct conformational responses. Instead of a complete transition from the up state to the down state in WT, both mutants exhibited a partial transition to an intermediate state (Fig. 5). A quick transition (<1 μs) to the intermediate state was consistently observed in three independent runs for R222Q (Fig. 6A), but the up-to-intermediate transition is much slower (>4 μs) in two runs for R811Q (Fig. 6B). Another simulation for R811Q was still in the up state at the end of 5μs. This suggests that the equivalent mutations from different VSDs can result in differential dynamic behavior of VSDs which may distinctly affect the functional outcomes.

**Figure 6.**
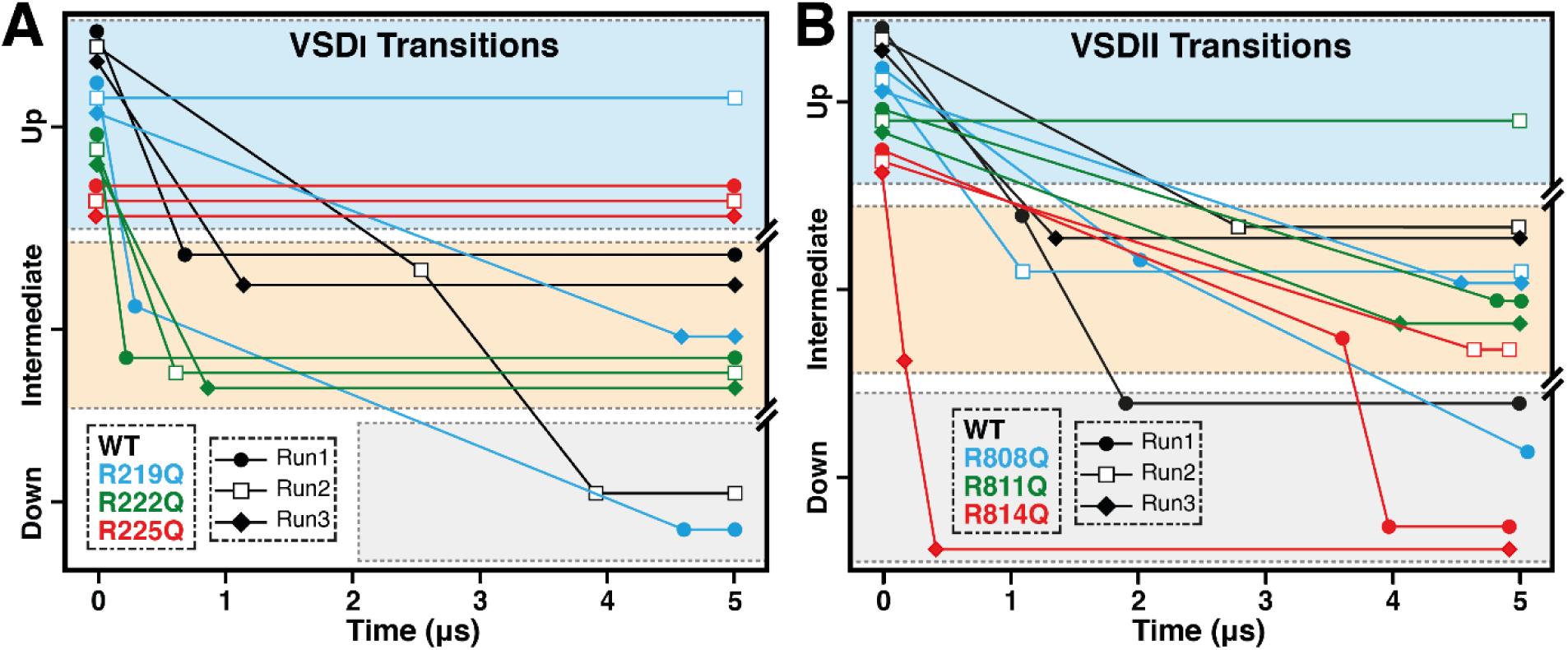
Comparative analysis of MD up-to-down transition rates for WT and R-to-Q mutations in VSD_I_ and VSD_II_. (A) Dynamic behaviors of VSD_I_ transitions, showing WT (black), R219Q (blue), R222Q (green), and R225Q (red). (B) Dynamic behaviors of VSD_II_ transitions, showing WT (black), R808Q (blue), R811Q (green), and R814Q (red). Data points represent results from three independent simulation runs, depicted by filled circles, empty squares, and filled squares. All simulations begin in the up state (light blue) under an applied external electric field of -500 mV, with transitions observed towards the intermediate state (orange) and down state (gray).

### R-to-Q mutations induce differential impacts on leaky gating-pore (I_ω_) current

It has been reported that several gating-charge mutants lead to gating-pore (I_ω_) currents leaking through the VSD (Chen et al., 1996; Daniel et al., 2019; Gosselin-Badaroudine et al., 2012b; Laurent et al., 2012; Mann et al., 2012; Moreau et al., 2018, 2015b, 2015a; Nair et al., 2012). To characterize the structural basis of this phenomenon, the minimum pore radius in the VSDs was analyzed through all trajectories and compared among the WT and all six mutants, which demonstrated distinctive pore opening between VSD_I_ and VSD_II_ (Fig. 7).

**Figure 7.**
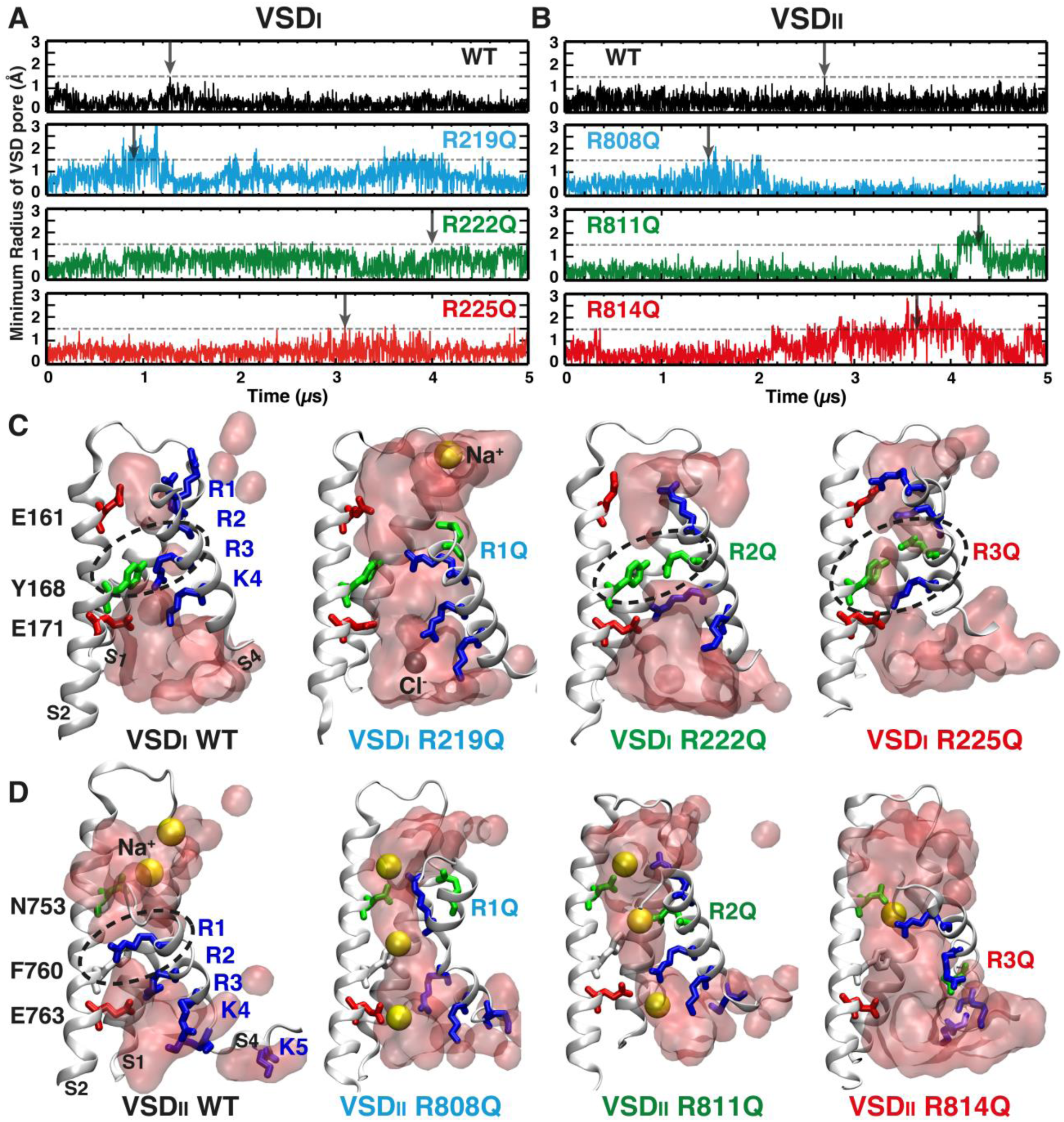
Pore opening of mutants in VSD_I_ and VSD_II_ during MD simulations. (A and B) Each panel depicts the time series of the minimum radius of the aqueous pathway of gating pore through VSDs during simulations. A 1.5 Å dashed line represents the potential pore opening threshold for water and ion permeation through VSDs. WT traces are presented in black, R1 mutants (R219Q and R808Q) in blue, R2 mutants (R222Q and R811Q) in green, and R3 mutants (R225Q and R814Q) in red. (C-D) Each panel shows molecular images of VSD_I_ (C) and VSD_II_ (D) WT and Mutants (S3 removed for clarity). In each snapshot, gating charges (blue), countercharges (red), HCS (green) residues are illustrated in licorice representation. Sodium (yellow) and chloride (gray) ions are depicted in spheres. Snapshots are selected at the time points indicated by black arrows in A and B. The water molecules are illustrated as a transparent red surface. The black dashed circles indicate the non-accessible solvent region, separating the extracellular and intracellular water milieu.

Remarkably, the minimum radius of the gating pore formed within VSD_I_ and VSD_II_ for WT while transitioning to the down state, remained below 1.5 Å throughout the simulation time (Fig. 7A&B) with HCS separating the intracellular and extracellular water crevices (Fig. 7C&D). This created a hydrophobic gate, which was not permeable to the gating-pore current. However, several equivalent gating-charge mutants exhibited differential permeability in simulations. For instance, the VSD structure of R222Q was not permeable throughout the entire trajectory, including the up state, the intermediate state, and the transitions between them (Fig. 7A). However, the equivalent R2 mutant in VSD_II_, R811Q, exhibited increased gating-pore opening during the intermediate state (Fig. 7B). This was characterized by a larger minimum pore radius exceeding 1.5 Å and the permeation of Na^+^ ions, as evidenced by the formation of a complete water wire (Fig. 7C&D) and multiple Na^+^ ion passage (Supp. Fig. 2B). Similarly, the R3 mutant in VSD_I_, R225Q, remained impermeable in its up state throughout the entire simulation, whereas the equivalent mutant R814Q exhibits gating pore with several Na^+^ ions permeating through.

A notable observed phenomenon was that gating-pore formation and Na^+^ ion permeation occurred concurrently with the structural transition. A correlation was observed between the time series for the minimum radius and the z-position of gating charges representing the state transition. Intermediate states of mutants were more likely to exhibit leakier gating-pore than WT as shown in (Supp. Fig. 2). Specifically, the gating-pore current was observed in VSD_II_ mutants —R808Q, R811Q, and R814Q— aligning with the state transitions at approximately 1.8 μs, 4.2 μs, and 3.9μs, respectively, as detailed in (Supp. Fig. 2B). Similarly, by tracking Na^+^ ions along z-axis, Ion permeation events were aligned with the minimum radius peak (Supp. Fig. 2).

### The state-dependent salt-bridge network explains the differential mutational effects

To elucidate the structural basis by which gating-charge mutations differentially influence the dynamics of the Na_v_ channel, key interactions within the VSD for each WT/mutant were analyzed across various channel states. This analysis unravels the intricate salt-bridge interactions between gating-charge, countercharge residues, and lipid molecules. State-dependent salt-bridge analysis was performed to quantify the occupancy between gating charge and countercharge residues, along with gating charge interactions with lipid molecules, across distinct structural states (up, intermediate, and down) (Fig. 8). Notably, our findings highlighted the state dependency and structural context as key points for understanding how the mutations altered the stability and dynamics of the salt-bridge interactions that modulate the voltage sensing and the gating-pore opening.

**Figure 8.**
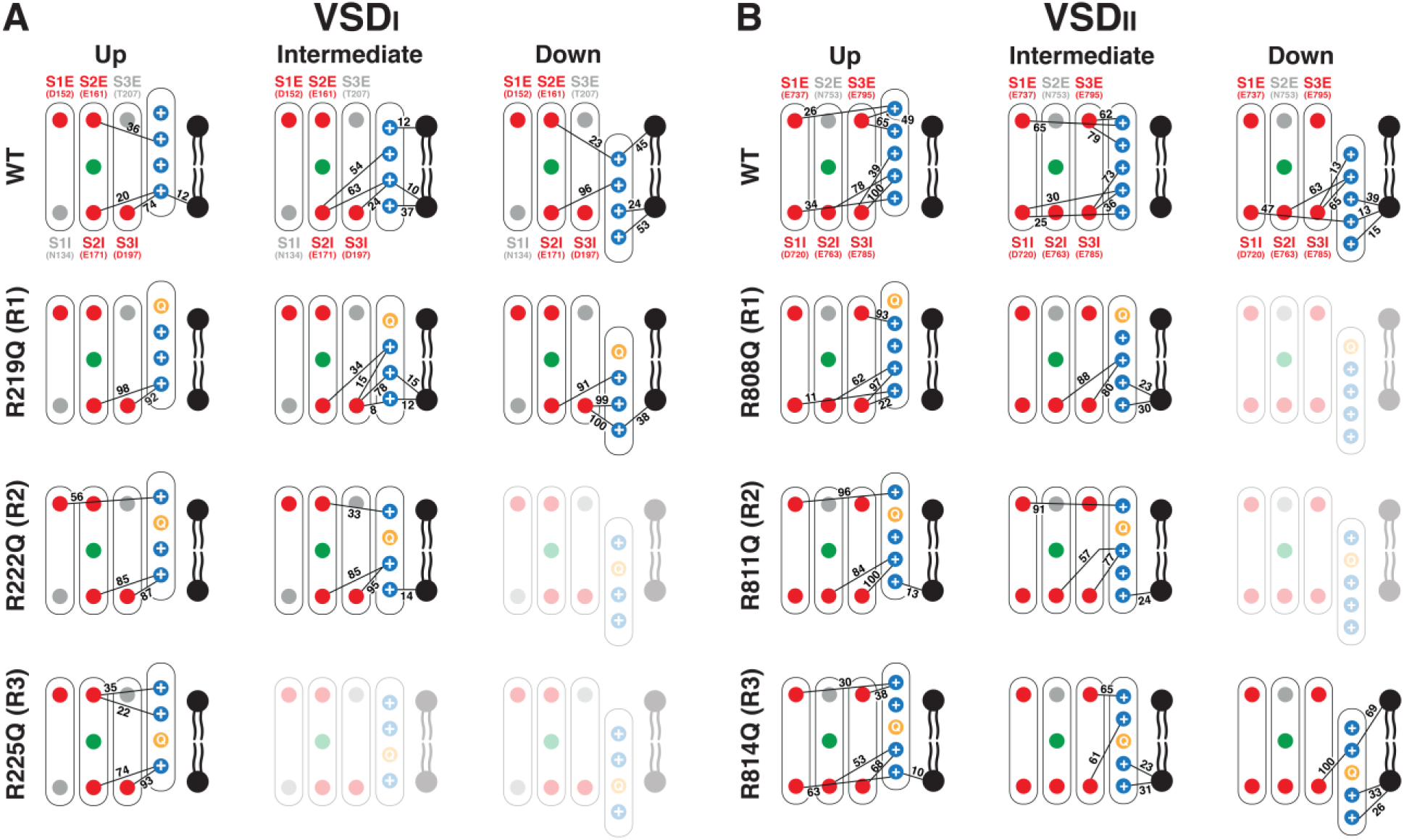
State-dependent salt-bridge network analysis for R-to-Q mutants in VSD_I_ and VSD_II_. Comprehensive analysis for WT and mutants (R1, R2, and R3) in VSD_I_ (A) and VSD_II_ (B). Each panel depicts the map of electrostatic interactions at various states (up, intermediate, and down) within each VSD. Nodes represent residues of gating charges (blue), mutated residues (orange), countercharges (red), HCS (green), and lipids (black). Edges represent the salt bridges, with numbers indicating the occupancy percentage of the interaction during each state. Countercharges are labeled with residue IDs from Na_v_1.5.

Upon comparing the salt-bridge networks of VSD_I_ and VSD_II_, VSD_II_ possessed a more intricate network compared to VSD_I_. This is due to the presence of a greater number of nodes—five gating-charge residues and five countercharges in VSD_II_ (Fig. 8B) compared to four gating charges and four countercharges in VSD_I_ (Fig. 8A). This complexity suggests variations in the stabilization mechanisms between the two VSD domains, as several critical hubs were identified in both. The state-dependent salt-bridge network analysis further highlighted key differences between WT of VSD_I_ and VSD_II_. In VSD_I_, the R3 gating charge served as an interaction hub in the intermediate state, interacting with countercharge INC residues (S2I-E171) and (S3I-D197) at occupancy of 63% and 24% respectively. However, R3 in VSD_II_ does not play a similar role as an interaction hub. Instead, it had only one interaction, with an occupancy of 73%, involving a countercharge (S3I-E785) in the intermediate (Fig. 8).

Salt-bridge provides insights into the contrasting dynamic behavior of R-to-Q mutations on VSD_I_ and VSD_II_. The R3 mutant in VSD_I_ (R225Q) exhibited resistance to the transition from the up state. This resistance arises from two primary factors: 1) the neutralization of the R3 gating charge residue disrupts multiple interactions with both countercharged INC residues and lipid molecules in the intermediate state and 2) this mutation does not destabilize the up state, as all salt bridges in the up state are maintained (Fig. 8A). As a result, the transition to the intermediate state becomes energetically unfavorable, explaining the observed resistance in all three replicas. In contrast, the R814Q mutation in VSD_II_ undergoes a complete transition to the down state. This difference stems from the fact that R3 in VSD_II_ forms only one salt bridge during all states, unlike its critical hub role in VSD_I_ (Fig. 8B).

The R2 mutants in VSD_I_ and VSD_II_ (R222Q and R811Q) both transitioned to the intermediate state during 5 µs simulations but exhibited different kinetic behaviors. All three independent runs of R222Q reached intermediate state within 1 µs (Fig. 6A). In contrast, R811Q took over 4 µs to transition to the intermediate state in two trajectories, while in another run, it maintained in the up state at the end of the trajectory (Fig. 6B). This difference can be attributed to the higher salt-bridge occupancies in the up state for R811 compared to R222Q. For instance, the occupancies of two salt bridges in R811Q (S3I-R4 and S1E-R1) were 100% and 96%, whereas the equivalent two in R222Q were 87% and 56%. It is reasonable that breaking two stronger salt bridges requires more time. Additionally, the salt-bridge network analysis also provided clues as to why both R222Q and R811Q did not reach the down state within 5 µs. In the WT, the R2 exhibits a high occupancy of salt bridges in both VSD_I_ and VSD_II_ in the down state (Fig. 8). The absence of such a crucial gating charge likely rendered the down state energetically unfavorable for the R2 mutants.

Furthermore, salt-bridge analysis explains the differing impacts of R-to-Q mutations on the leaky gating-pore current between VSD_I_ and VSD_II_ mutants. While R2 and R3 of VSD_II_ (R811Q and R814Q) exhibited a gating-pore opening during transitions, R2 and R3 of VSD_I_ (R222Q and R225Q) did not display such a gating pore in any of the replicas. This difference can be attributed to the lower occupancy of salt bridges in the intermediate state of VSD_II_ compared to VSD_I_. Specifically, In VSD_I_, R222Q demonstrated a higher interaction occupancy between gating charges and INC countercharges residues (S2I-E171) and (S3I-D197) at occupancies of 85% and 95% respectively (Fig. 8A), compared to 57% and 77% in VSD_II_ (Fig. 8B). This higher occupancy helps maintain VSD_I_ in a compact form at the INC compared to VSD_II_, explaining the absence of leakage during transition. Additionally, VSD_II_ is more hydrophilic as it contains complete or partially unoccupied charged residues, facilitating ions’ passage through the VSD gating pore and into the cell. In summary, the differential impacts of R-to-Q mutations on gating-pore currents can be attributed to variations in the interaction occupancies and availability of countercharge residues, leading to differences in the hydrophilicity and polarity of the domains.

## Discussion

The analysis of disease-associated missense variants highlighted gating charges in the S4 helix of VSD as major mutational hotspots, emphasizing their crucial role in Na_v_ functionality. Previous electrophysiological recordings have shown that in the WT channel, the S4 segments rapidly return to their resting conformation after repolarization (Albert et al., 1999; Gamal El-Din et al., 2014). In contrast, studies have reported that the mutated S4 segments remain trapped in conductive (activated) conformations even at hyperpolarized voltages, consistent with the immobilization of the S4 segment that has been proposed to underlie the formation of gating pore currents(Moreau et al., 2015b). Employing MD simulations, coupled with controlled external electric fields (±500 mV), allowed for the study of structural transitions in VSDs, providing insights into the conformational responses of both WT and mutant VSDs. Notably, mutations (R-to-Q) in equivalent gating-charge positions from different VSDs displayed distinct dynamic behaviors during the transition from the up state to the down state, as well as differential leaky I_ω_. Analyzing the state-dependent salt-bridge network for WT and mutant trajectories reveals molecular mechanisms behind these differential mutational effects.

Gating charges and countercharges within each VSD are highly conserved within the Na_v_ family, as demonstrated by the MSA of nine human Na_v_ channels (Fig. 2). This conservation suggests that homologous gating-charge/countercharge mutations, such as the R2 mutants in VSD_II_ in both Na_v_1.4 and Na_v_1.5, may have similar effects across different Na_v_ isoforms/subtypes, as supported by our recent study showing an 86% agreement in gating properties among homologous variants across various Na_v_ channels (Ahangar et al., 2024). However, MSA reveals distinct numbers and distributions of gating charges and countercharges for each VSD domain (Fig. 2). The count and locations of these charges vary across VSD domains, giving each VSD a unique salt-bridge network and distinct structural environment for every gating charge. Therefore, mutational impacts in VSD_III_ and VSD_IV_ cannot be inferred from their equivalents in VSD_I_ and VSD_II_ (Albert et al., 1999), highlighting the complexity of VSD structures and the need for further research to understand the structural interplay in VSD_III_ and VSD_IV_ fully.

While the countercharges S2I and S3I are highly conserved, the other four countercharge positions (S1I, S1E, S2E, and S3E) show variability among VSDs (Fig. 2). The conservation of S2I and S3I underscores their functional importance in Nav channels. The significance of S3I is further corroborated by its identification as a mutational hotspot (Fig. 3). The remaining four countercharge positions exhibit conservation only within specific VSDs, rather than across all four VSDs. For example, S3E is conserved as glutamate solely in VSD_II_ among the nine Na_v_ isoforms, while the equivalent position is occupied by T/S/G in VSD_I_, VSD_III_, and VSD_IV_ (Fig. 2). Similarly, S2E is conserved as acidic residues only in VSD_I_ and VSD_III_, but is replaced by asparagine in the other VSDs (Fig. 2). As mapped in the salt-bridge networks (Fig.8), this VSD- specific distribution pattern of countercharges contributes significantly to the differential impacts of gating-charge mutations.

Interactions between lipids and gating charges form an integral part of the salt-bridge network in VSDs. This suggests that the composition of lipids can significantly influence the kinetics of state transitions, which agrees with the previous studies on lipid regulation on Na_v_ channels (Bendahhou et al., 1997; Kang et al., 1996; Wieland et al., 1996). Different types of lipids can interact distinctly with gating charges, potentially affecting the stability of each state and the transition rates between states(D’Avanzo et al., 2013). Furthermore, the unique structural characteristics of each VSD can influence the accessibility of gating charges to lipids. Each VSD domain has a distinct number of gating and countercharges, which can affect how these residues interact with surrounding lipids (Sands and Sansom, 2007; Schmidt et al., 2006; Zheng et al., 2011). For instance, the R1 gating charge (R1599) in VSD_IV_ of Na_v_1.7 showed ion-pair interaction with the phosphodiester group of a POPC lipid (Ahuja et al., 2015). These gating charges engage in compensatory interactions with phospholipids, thereby stabilizing different gating states of the VSD and channel (Schmidt et al., 2006; Xu et al., 2008). This protein-lipid interaction could influence the stability and transition kinetics of each state. Further studies could provide more insights into the influence of membrane lipids on eukaryotic Na_v_ channel activity and their implications for the function of VSDs.

Differential gating-pore openings caused by gating-charge mutations are observed between VSD_I_ and VSD_II_, attributed to the distinct characteristics of these domains. Specifically, equivalent mutations in VSD_II_ are more permeable than those in VSD_I_. This increased permeability in VSD_II_ is likely due to more countercharge residues than VSD_I_, which facilitates ion permeability through the gating pore. Additionally, the higher occupancy of salt bridges in the intermediate state of R222Q mutant in VSD_I_ makes the system more compact at the INC, reducing its openness and permeability. Both R219Q and R808Q showed gating pore opening during the intermediate state (supp. Fig. 2). This phenomenon can be attributed to a reduction in the interaction occupancy between gating charges and countercharged residues in the transition from up to intermediate states (Fig. 8). The R219Q findings are consistent with previously reported experimental data (Moreau et al., 2018). However, for R808Q, existing experimental data lacks measurements of I_ω_, precluding direct comparison with our observations (Chen et al., 1996; Glazer et al., 2020).

It is important to acknowledge several limitations in our computational study, despite performing over 120 μs of simulations in total. Firstly, although each simulation ran for 5 μs, this timescale proved insufficient to capture the complete transitions of some mutants such as R222Q, R225Q, and R811Q (Fig. 6). Extending the simulation time much longer (> 20 µs) and conducting more independent runs would likely to capture the “up” to “down” transition and provide a more statistically robust characterization of each mutant’s behavior. However, such an approach requires computational resources beyond our current capacity. The missing “down” state explains the discrepancies between our simulations and experimental data in gating-pore permeability, while experiments showed ion permeability for R222Q mutations in the resting (“down”) state (Daniel et al., 2019; Moreau et al., 2015b), we did not capture this phenomenon, likely due to the absence of the down (resting) state for R222Q in the simulations (Francis et al., 2011; Jiang et al., 2018; Sokolov et al., 2007). Secondly, our simulations were limited to the up-to-down transitions of VSD_I_ and VSD_II_, representing only two sub-steps of the recovery process in Na_v_ channels (Ahern et al., 2016; Albert et al., 1999; Goldschen-Ohm et al., 2013; Lacroix et al., 2013). This limitation impedes the direct comparison between our computational results and electrophysiological measurement of recovery rate. Thirdly, our study focused exclusively on the VSDs and did not include the potential effects of the mutations on the coupling between the VSDs and the pore domain (Chanda et al., 2004; Chowdhury and Chanda, 2012; Cowgill and Chanda, 2021; Muroi et al., 2010). To fully understand the functional implications of these mutations, simulations of the entire channel, including the pore domain and the interactions between the VSDs and the pore, would be the next step of our following study. Overall, to achieve a comprehensive understanding of the impacts of these gating-charge mutations on the functional cycle and to enable a comparison with all gating properties measured in electrophysiology, the mutations need to be introduced into a full-length model of a Na_v_ channel, and simulations should be performed for all state transitions. Such a comprehensive study was not achieved in the present study due to limited computing resources, but it remains a long-term objective for our future research.

## Acknowledgments

Computer resources came from a Maximize ACCESS allocation through project BIO210015, an allocation (MCB200085P) on Antons at the Pittsburgh Supercomputing Center provided by the National Center for Multiscale Modeling of Biological Systems through National Institutes of Health grant P41GM103712-1 and from a loan from D. E. Shaw Research, and a Frontera Pathways allocation (MCB21012) at the Texas Advanced Computing Center (TACC). Research reported in this publication was supported by an Institutional Development Award (IDeA) from the National Institute of General Medical Sciences of the National Institutes of Health under award number P20GM130460, the Data Science/AI Research Seed Grant (SB3002 IDS RSG- 03) from Institute for Data Science at the University of Mississippi.

## Author Contributions

Conceptualization: EE, AA, TGE, JB, JL. Methodology: EE, AA, RR, JB, JL. Investigation: EE, AA, RR. Visualization: EE. Supervision: JL. Writing—original draft: EE, JB, JL. Writing—review & editing: EE, AA, RR, TGE, JB, JL

## Declaration of Generative AI and AI-assisted technologies in the writing process

During the preparation of this work, the author(s) used ChatGPT to assist with polishing the text. After using this tool/service, the author(s) reviewed and edited the content as needed and take(s) full responsibility for the content of the publication.

## Supplementary Materials

### Materials and methods

#### Plasmids and Mutagenesis

The plasmid from human Na_v_1.5 (hNa_v_1.5) was a generous gift from Peter Ruben (Simon Fraser University). GFP plasmid used for co-expression with hNa_v_1.5 plasmid was purchased from Lonza. The hNa_v_1.5 mutants used in this study were made using site-directed mutagenesis in- house and verified using whole plasmid sequencing services through Plasmidsaurus.

#### Cell Culture and Transfection

CHO-K1 cells (ATCC) were cultured using Ham’s F12 medium with 10% FBS at 37°C in 5% CO_2_. Cells were grown to ∼70-80% confluency and were co-transfected with wildtype (WT) or mutant human Na_v_1.5 plasmid and GFP plasmid DNA via electroporation with a Lonza 4D Nucleofector unit following the manufacturer’s protocols. Following transfection, cells were plated on 12-mm glass coverslips coated in poly-L-lysine and incubated at 30°C in 5% CO_2_.

#### Electrophysiological Recordings

All experiments were performed 16–30 h after transfection at room temperature. A whole-cell patch-clamp configuration was used to assess peak current magnitudes. Borosilicate glass pipettes (Harvard Apparatus) were pulled to a resistance of 2–6 MΩ (P-1000; Sutter Instrument). Glass pipettes were filled with an internal solution containing (in mM) 130 CsF, 10 NaF, 10 EGTA, and 10 HEPES, pH 7.4 adjusted with CsOH. The extracellular solution contained (in mM) 50 NaCl, 100 NMDG, 10 HEPES, 2 CaCl_2_, and 1.8 MgCl_2_, pH 7.4 adjusted with NaOH. An Axopatch 200B amplifier and pCLAMP 10.6 (Axon Instruments) were used to record whole-cell currents. All recordings were performed with a starting holding potential of −80 mV with a 5-kHz low-pass filter and sampling at 10 kHz. The series resistance measured in whole-cell configuration was compensated by 70-80%. Fluorescence was visualized on an Olympus IX73 microscope with a CoolLED pE-4000 illumination system. Only cells that were fluorescent for GFP and produced hNa_v_1.5 currents were analyzed in electrophysiological experiments.

To measure the voltage dependence of steady-state activation, currents were elicited using a voltage-clamp protocol where depolarizing pulses were applied for 100 ms from -90 to 70 mV in 10 mV increments. Peak currents were analyzed using a custom-written MATLAB script and activation curves were then generated using a standard Boltzmann equation. To measure the voltage dependence of steady-state inactivation, a conditioning prepulse was applied to membrane potentials ranging from a holding potential of -130 to -10 mV for 200 ms in 10 mV increments before measuring non-inactivated channels using a 100 ms pulse to -10 mV at each step. The currents elicited following the preconditioning pulse were then analyzed using a custom-written MATLAB script and inactivation curves were fitted to a standard Boltzmann equation. To measure the recovery from inactivation, currents were elicited using a two-pulse protocol at 10 mV to obtain maximal activation. The second pulse was elicited at varying time points starting at 1 ms and increasing to 1024 ms doubling every increment. Channel recovery was determined by normalizing the current elicited from the second test pulse to the first conditioning pulse and plotted against the recovery time. The curves were fitted with a single exponential function.

#### Data Analysis

Data were analyzed using pCLAMP 10.6 (Axon Instruments) and custom-written MATLAB programs (The MathWorks Inc.). Data are expressed as means ± SEM. Data were compared using a one-way ANOVA with a post-hoc Dunnett’s test.

### Results

#### Equivalent mutations show differential gating-property impacts in electrophysiology

While many of these mutations have been examined functionally in previous studies, they have never been directly compared in the same system and the same study. Some of the measured gating differences could be due to differences in experimental setup including, solutions, cell system, or the presence or absence of accessory subunits. To show that identical mutations in equivalent arginine positions can cause different changes to channel gating, we compared a number of channel properties at the R2 position of VSD_I_ and VSD_II._ We used whole cell patch clamp measurements in Chinese Hamster Ovary (CHO) cells transfected with Na_V_1.5 from humans. First, we determined the voltage dependence of activation for WT Na_V_1.5 along with R-to-Q mutations at R2 in VSD_I_ and VSD_II_. Supplementary Figure 1A shows representative current families from each channel. In the R2 position, mutations to glutamine have opposite effects in VSD_I_ and VSD_II_ with R222Q showing a hyperpolarizing shift in the voltage-dependence of activation. In contrast, R811Q shows a depolarizing shift (Supp. Fig. 1B). Contrastingly, mutations in VSD_II_ show minimal changes in the voltage dependence of activation. We then measured the voltage-dependence of inactivation and found that R222Q and R811Q mutations show a trend towards a hyperpolarizing shift with significant shifts in R222Q in VSD_I_ (Supp. Fig. 1C). Finally, we found that recovery from inactivation was slowed in only the R811Q mutation but WT-like in the R222Q (Supp. Fig. 1D). Taken together, these results suggest that identical mutations in equivalent positions in different VSDs can have complex and unpredictable changes in channel function. Our goal is to use computational approaches to begin to look at how these mutations alter channel structure and dynamics with the hopes of building models where we can begin to predict functional changes based on these structural changes.

---

**Supplementary Figure 1:**
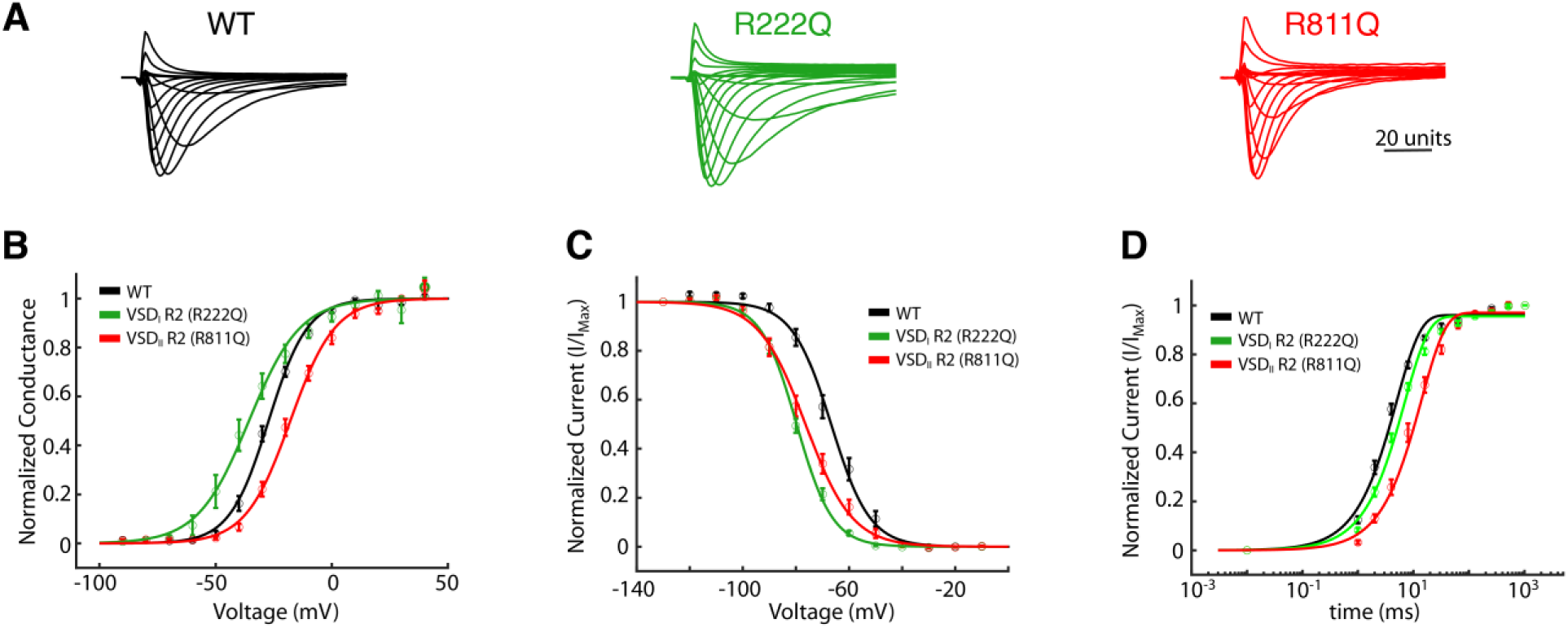
Functional measurements for mutations in VDS_I_ and VSD_II_ R2 to glutamine. (A) Representative currents for each channel in response to a series of 100ms voltage pulses from -90 to 70 mV in 10 mV increments. (B) Conductance versus voltage plots comparing WT (n=9) to R222Q (n=5) and R811Q (n=6). (C) Steady-state channel availability curves comparing WT (n=8) to R222Q (n=7) and R811Q (n=7). Plots showing the recovery time from inactivation again comparing WT (n=7) to R222Q (n=7) and R811Q (n=8). Fits were performed as described in the methods and the results are shown in Supp.Table 2 along with the determination of significance using a one-way Anova with a post-hoc Dunnett’s test.

**Supplementary Figure 2:**
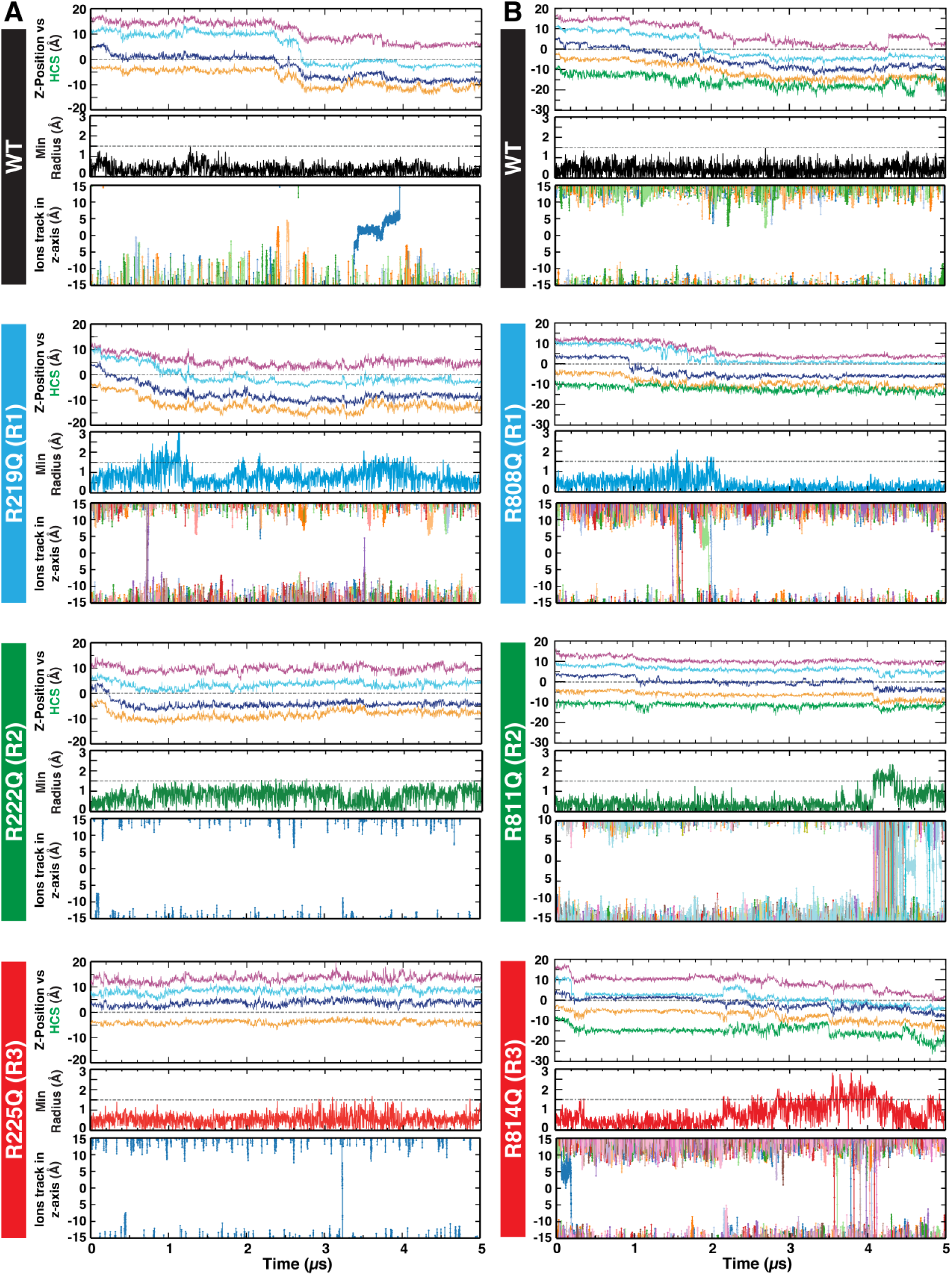
**The time series of VSD structural transition, gating pore opening, and ion permeations**. This figure presents a comprehensive study of WT and mutants (R1, R2, R3) in VSD_I_ (A) and VSD_II_ (B). Each panel shows the z positions of gating charge residues relative to HCS (top), the minimum pore radius of VSDs (middle), and ion permeations (bottom) with the function of time.

**Supplementary Table 1:** Summary of MD system composition, simulation durations, and the number of replica simulations for both WT and mutant systems.

| System | Composition | # Atoms | Duration (μs) | # of Replicas |
| --- | --- | --- | --- | --- |
| WT (VSD<sub>I</sub>) | VSD<sub>I</sub> (119 - 250) PD<sub>II</sub> (840 - 944)<br>123 POPC and 41 POPI<br>8651 Water Molecules<br>NaCl [60 Na<sup>+</sup> and 22 Cl<sup>-</sup>] | 52012 | 5 | 3 |
| R219Q | VSD<sub>I</sub> (119 - 250) PD<sub>II</sub> (840 - 944)<br>123 POPC and 41 POPI<br>8651 Water Molecules<br>NaCl [61 Na<sup>+</sup> and 22 Cl<sup>-</sup>] | 52005 | 5 | 3 |
| R222Q | VSD<sub>I</sub> (119 - 250) PD<sub>II</sub> (840 - 944)<br>123 POPC and 41 POPI<br>8651 Water Molecules<br>NaCl [61 Na<sup>+</sup> and 22 Cl<sup>-</sup>] | 52005 | 5 | 3 |
| R225Q | VSD<sub>I</sub> (119 - 250) PD<sub>II</sub> (840 - 944)<br>123 POPC and 41 POPI<br>8651 Water Molecules<br>NaCl [61 Na<sup>+</sup> and 22 Cl<sup>-</sup>] | 52005 | 5 | 3 |
| WT (VSD<sub>II</sub>) | VSD<sub>I</sub> (699 - 838) PD<sub>II</sub> (1329 - 1480)<br>99 POPC and 33 POPI<br>9114 Water Molecules<br>NaCl [58 Na<sup>+</sup> and 22 Cl<sup>-</sup>] | 49954 | 5 | 3 |
| R808Q | VSD<sub>I</sub> (699 - 838) PD<sub>II</sub> (1329 - 1480)<br>99 POPC and 33 POPI<br>9114 Water Molecules<br>NaCl [58 Na<sup>+</sup> and 23 Cl<sup>-</sup>] | 49946 | 5 | 3 |
| R811Q | VSD<sub>I</sub> (699 - 838) PD<sub>II</sub> (1329 - 1480)<br>99 POPC and 33 POPI<br>9114 Water Molecules<br>NaCl [58 Na<sup>+</sup> and 23 Cl<sup>-</sup>] | 49946 | 5 | 3 |
| R814Q | VSD<sub>I</sub> (699 - 838) PD<sub>II</sub> (1329 - 1480)<br>99 POPC and 33 POPI<br>9114 Water Molecules<br>NaCl [58 Na<sup>+</sup> and 23 Cl<sup>-</sup>] | 49946 | 5 | 3 |

**Supplementary Table 2:** Summary of gating properties of mutations. . Voltages of half-activation and inactivation were calculated from the average of fits, using the Boltzmann function, to the experimental data from each cell. Tau of recovery was calculated as the average of fits to a single exponential function. Values are reported as the mean ± S.E. Statistical significance from a one-way ANOVA with a post-hoc Dunnet’s test is indicated as follows: * = p<0.05, ** = p<0.01, *** = p<0.001, **** = p<0.0001.

| hNa<sub>v</sub>1.5 WT Channel | Voltage-dependence of activation |  |  | Closed-state Inactivation |  |  | Recovery from Inactivation |  |  |
| --- | --- | --- | --- | --- | --- | --- | --- | --- | --- |
| | $V_{1/2}$ (mV) | $k$ | $n$ | $V_{1/2}$ (mV) | $k$ | $n$ | $\tau$ (ms) | Weight (%) | $n$ |
| WT | -27.25 ± 1.06 | -8.63 ± 0.36 | 9 | -66.80 ± 1.73 | 7.23 ± 0.17 | 8 | 4.98 ± 0.34 | 96.06 ± 0.69 | 7 |
| R222Q (VSD1) | -35.71 ± 3.16* | -11.08 ± 0.61** | 5 | -79.73 ± 0.93**** | 6.71 ± 0.30 | 7 | 6.99 ± 0.29 | 95.60 ± 0.58 | 7 |
| R811Q (VSD2) | -17.89 ± 1.51* | -9.70 ± 0.22 | 6 | -76.41 ± 1.71*** | 8.60 ± 0.35* | 7 | 14.00 ± 1.41**** | 97.06 ± 0.69 | 8 |

